# AI-based prediction of herbarium sequencing success across the plant tree of life

**DOI:** 10.1101/2025.02.03.636220

**Authors:** Yasaman Ranjbaran, Olivier Maurin, Elena Canadelli, Tomas Morosinotto, Marie-Helen Weech, Paul Kersey, Alexandre Antonelli, William J. Baker, Gabriele Sales, Francesco Dal Grande

## Abstract

DNA recovered from herbarium specimens represents a vital asset in botanical research, playing a pivotal role in unravelling the evolution, diversity, and ecological dynamics of plants. Despite its importance, challenges such as fragmented DNA and insufficient sequencing yields render molecular data retrieval a high-risk and costly endeavour involving the use of non-replaceable herbarium specimens. Here, we propose a framework based on Artificial Intelligence (AI) to forecast the success of genomic DNA extraction suitable for sequencing from herbarium samples. Our model integrates morphological characteristics and sample colour derived from scanned herbarium images, metadata including sample age and locality, and DNA quantity measurements of samples. We train a deep learning algorithm with ca. 2,000 specimens that have been digitized and sequenced in the framework of the Plant and Fungal Trees of Life (PAFTOL) Project, spanning from year 1832 to the present. As training datasets increase with ongoing digitization and genomic sequencing efforts, our AI predictive model can support researchers in selecting the herbarium samples with the highest likelihood of yielding high-quality genomic DNA from amongst a vast array of globally distributed candidate specimens. Our approach enhances the contribution of herbarium-derived DNA in large-scale studies and facilitates the utilisation of historical collections for a deeper understanding of plant evolution and ecology, with implications for conservation.

## Introduction

Biodiversity is facing unprecedented global decline, driven by climate change, habitat destruction, and other human-induced pressures (Diaz et al., 2019; IPES, 2022; WWF, 2020). This accelerating loss in biodiversity and ecosystem degradation diminishes the pool of species and lineages that sustain essential ecosystem services (e.g., food security, water purification, and climate regulation), disrupts the ecological and evolutionary processes essential for sustaining the long-term resilience of ecosystems (Adla et al., 2022; Xu et al., 2020) and is now considered the second-most-severe risk facing humanity in the next decade (WEF, 2025).

### Genomics as a Tool for Conservation

To mitigate these impacts, conservation strategies must be informed by a robust understanding of not only the diversity and distribution of species (Antonelli et al., 2023), but also the evolutionary processes that underpin their resilience to environmental change (Pearman et al., 2024). Fortunately, genomic research has transformed and accelerated our capacity to investigate such processes. High-throughput sequencing and advanced analytical frameworks now provide unprecedented insights into the genetic mechanisms by which organisms respond to stressors (Lancaster et al., 2022; Waldvogel et al., 2020; Bernatchez et al., 2024). Key insights include detecting shifts in allele frequencies (Frankham, 2010; Hohenlohe et al., 2021; Moore et al., 2021), tracking effective population size over time (Hayes et al., 2003; Roman & Palumbi, 2003), and identifying genetic erosion (Díez-del-Molino et al., 2018; Gargiulo et al., 2024). Together, these findings clarify how species adapt and survive under novel conditions (Wisser et al., 2019). These insights not only deepen our understanding of evolutionary dynamics but also enable the identification of species and populations most at risk (Hohenlohe et al., 2021).

Plants account for a disproportionately large share of threatened species worldwide (IUCN, 2025). Among species classified as Critically Endangered, Endangered, or Vulnerable by the International Union for Conservation of Nature Red List of Threatened Species, 28,159 (61%) are plants (IUCN, 2024), and about 45% of all flowering plants are now estimated to be threatened (Bachman et al., 2024). Plant diversity loss undermines ecosystem stability, disrupts nutrient cycling, and weakens resilience to environmental disturbances (Isbell et al., 2011). Safeguarding plant lineages is therefore critical and requires a robust understanding of their evolutionary potential and genetic variation. Yet, focusing on contemporary populations alone provides only a narrow snapshot of genetic diversity (Pääbo et al., 2004; Crisp et al., 2016).

### The Role of Herbarium Specimens and Artificial Intelligence (AI) in Conservation

Herbarium genomics offers unprecedented insights into population and phylogenomic studies (Roycroft et al., 2022) and can reveal how species have responded to environmental change over time, thereby enriching our ability to forecast and manage future threats (Shapiro and Hofreiter, 2014; Hofreiter et al., 2015; Johnoson et al., 2023) and produce information that could be incorporated into conservation policies (Jensen et al., 2023). Building on the unique temporal perspective that herbarium specimens provide, these centuries-old collections offer an unparalleled window into historical plant biodiversity (Lister, 2011; Bieker and Martin, 2018; Bakker et al., 2020). By bridging past and present populations, they enable researchers to trace genetic changes over time, detect signals of adaptation or genetic erosion, and reconstruct plant evolutionary histories (Hart et al., 2016; Meineke et al., 2018; Davis, 2024). For example, comparing 200-year-old specimens with their modern counterparts can reveal how genetic diversity has shifted in response to historical climate events (Papalini et al., 2023).

Herbarium specimens hold a remarkable potential to generate knowledge, but also present unique challenges as subjects of genomic research. DNA in these samples is often fragmented and chemically degraded due to enzymatic activity and extended storage, while microbial contamination further hinders recovery of high-quality data. Not every specimen yields sufficient genetic material for sequencing (Staats et al., 2013; Brewer et al. 2019; Kates et al., 2021; Papalini et al., 2023). These obstacles underscore the need for careful, strategic specimen selection to preserve non-renewable collections and maximize research benefits. Poorly planned sampling can waste resources and needlessly damage irreplaceable materials (Shepher, 2017; Davis et al., 2024).

Despite these challenges, recent breakthroughs in ancient DNA extraction and high-throughput sequencing have propelled herbarium genomics to the forefront (Bieker and Martin, 2018; Zeng et al., 2018; Bakker et al., 2020; Burbano and Gutaker, 2023). Large-scale initiatives such as the Plant and Fungal Trees of Life (PAFTOL) project coordinated by the Royal Botanic Gardens, Kew (RBG Kew) are now uncovering the evolutionary history of all plant and fungal genera and filling critical gaps in our understanding of the tree of life. However, PAFTOL relies on herbarium specimens for over one-third of their data, which exemplifies the transformative potential of herbarium-based research (Baker et al., 2022; Zuntini et al., 2024).

The mass digitization of herbarium collections has unlocked unprecedented opportunities to integrate genomic data with metadata (e.g., collection dates, geographic origins), thereby enriching evolutionary and ecological analyses (Soltis, 2017; Thiers et al., 2016; Smedt et al., 2024; Soltis et al., 2018). Over the past two decades, digitization efforts have expanded at institutional, national, and international scales (Nelson and Ellis, 2019), unlocking biodiversity data from an estimated 396 million specimens across 3,500 herbaria (Thiers, 2021). Platforms such as the Global Biodiversity Information Facility (GBIF) (2024), Integrated Digitized Biocollections (iDigBi) (2024), and the Reflora Virtual Herbarium (Reflora - Virtual Herbarium, 2024) now disseminate these data globally, and in 2019 alone, more than two studies per day cited GBIF resources (Paton et al., 2020). Yet a critical bottleneck remains; identifying which specimens are most likely to yield high-quality genomic data. Traditional heuristic criteria, such as specimen age and colour, often fail to predict sequencing success (Erkens et al., 2008), prompting calls for data-driven tools to optimize specimen selection (Bakker et al., 2016; Kates et al., 2021). Emerging technologies in AI, particularly deep learning (a type of machine learning that uses deep neural networks to learn patterns from data) (LeCun et al., 2015), offer promising solutions. By excelling in image-based tasks, deep learning-based models can autonomously detect patterns in digitized herbarium images, mitigating the limitations of manual feature extraction (LeCun et al., 2015).

Using assets generated by the PAFTOL project – encompassing a rich dataset of digitized herbarium samples, target capture sequence data, and associated metadata (Baker et al., 2023; Zuntini et al., 2024) – we develop a novel machine-learning tool to predict the DNA yield for sequencing from herbarium specimens. By autonomously identifying patterns linked to successful sequencing outcomes, our model pinpoints which specimens are best suited for genomic analysis and which should remain undisturbed. This data-driven approach has the potential to reduce costs, minimize destructive sampling (Davis et al., 2024), and maximize the long-term value of irreplaceable collections. Our findings highlight how AI can contribute to our scientific exploration of biodiversity and support responsible scientific practices.

## Materials and Methods

### Dataset Construction and Metadata Preprocessing

Metadata for 2,583 herbarium samples used in the PAFTOL project – 2,303 from the Kew Herbarium (Kew Data Portal, 2025) and 280 from the Naturalis Herbarium (Collections | Naturalis, 2025) – were initially recorded manually in tabular form. These samples are linked to genomic data generated under the PAFTOL project (Baker et al. 2022; Kew Tree of Life Explorer Data Release 3.0 [April 2023]). Records that lacked a value for the target variable, DNA concentration, were excluded from analysis, leaving 2,334 samples representing 1,481 species and 23 orders from 164 countries. Missing values in non-target variables were retained for later processing: for instance, 5% of the samples lacked an associated age, 21% a species indication, and 12% a country of origin.

Taxonomic annotations were initially assigned at the species level, but to capture the hierarchical structure of the taxonomic tree and mitigate the issue of many species being represented by a single sample, we extended these annotations to include the corresponding order and clade using the Taxoniq (https://github.com/taxoniq/taxoniq) and Taxize (https://f1000research.com/articles/2-191/v2) packages. Geographical data was supplemented with latitude and longitude coordinates for each country, enabling the capture of spatial relationships while retaining the original categorical information. To prevent overfitting, we identified any remaining factors with levels corresponding to a single observation and masked those with a missing value marker, thereby avoiding potential biases from underrepresented groups.

The original year of specimen collection was transformed into “sample age” by subtracting the collection year from 2024, the year of analysis. This distribution was long-tailed, so a logarithmic transformation followed by re-centring was applied to produce a more symmetric distribution. DNA concentration, measured in nanograms per microliter (ng/μL), showed a prominent peak at low values (∼10 ng/μL) followed by an exponentially decaying tail. A Boltzmann (Truncated Discrete Exponential) model was used to characterize this skew, after which a quantile transformation (SciPy library, version 1.14.1; 10.1038/s41592-019-0686-2) was employed to stabilize variance and improve interpretability.

### Image Segmentation

Segmentation of herbarium samples was performed using a convolutional neural network (CNN) built on the OCRNet architecture, implemented via the PaddleSeg framework (Liu et al., 2021). We trained this model using an external dataset published by White et al. (2020) which included 400 high-resolution fern images, each with a corresponding mask generated through a combination of automated and manual curation tools. Data augmentation techniques such as random rescaling, cropping, and rotation were employed to improve model robustness. In this procedure, 360 images were used for training and 40 for independent testing.

To enhance the quality of our training data, we removed herbarium images containing more than 98% blank pixels, as these images would contribute little to the training process. Additionally, we cropped the remaining images to remove extraneous elements, including white borders, dust specks, and small fragments detached from the samples (Figure 1).

**Figure 1.**
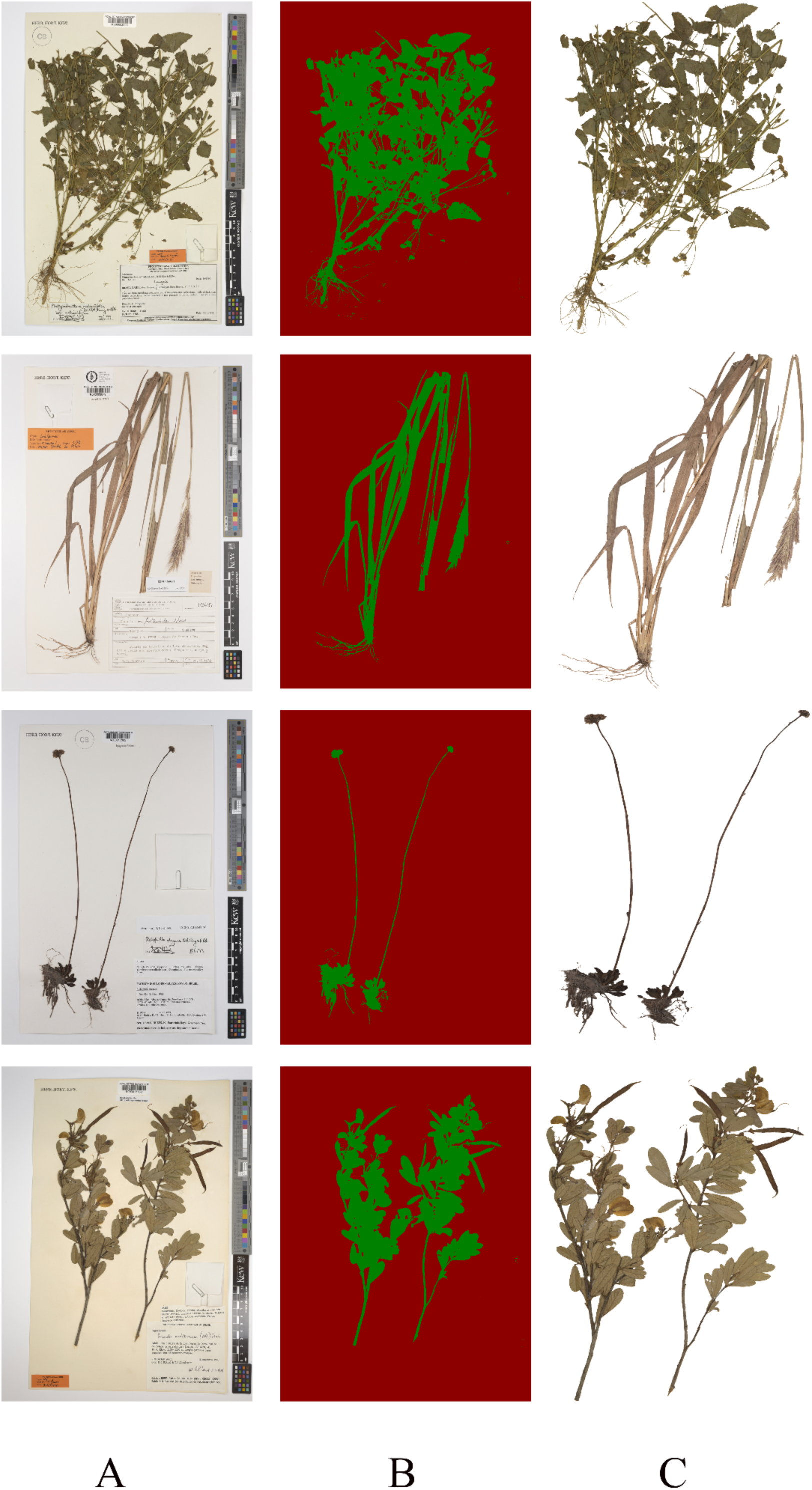
Three-step image segmentation process applied to four herbarium specimens, representing different plant shapes. (A) Original digitized herbarium sheets, including plant tissues, voucher information, and colour calibration labels. (B) Segmentation results showing the algorithm’s selection of plant tissues (highlighted in green) against the background. (C) Final output isolating only the plant tissues, with all non-plant elements removed. The algorithm successfully identified and separated plant tissues in different plant shapes, demonstrating its versatility and accuracy.

### Network Architecture

A multi-modal neural network was designed to predict DNA concentration from both herbarium images and tabular metadata (Figure 2). The image-processing module consists of a convolutional subnetwork based on Res2Net operating on 224 × 224-pixel images. We obtained pre-trained weights from the PyTorch Image Models library (Wightman, 2019; model ID: “timm/res2net50d.in1k”). To account for varying resolutions of individual herbarium sheets, each image is cropped to an 896-pixel square region, which is then divided into non-overlapping tiles. The CNN independently processed these tiles, generating 2,048-dimensional feature maps. These maps were then projected onto a 512-dimensional space through a learnable linear transformation, serving as a form of regularization.

**Figure 2.**
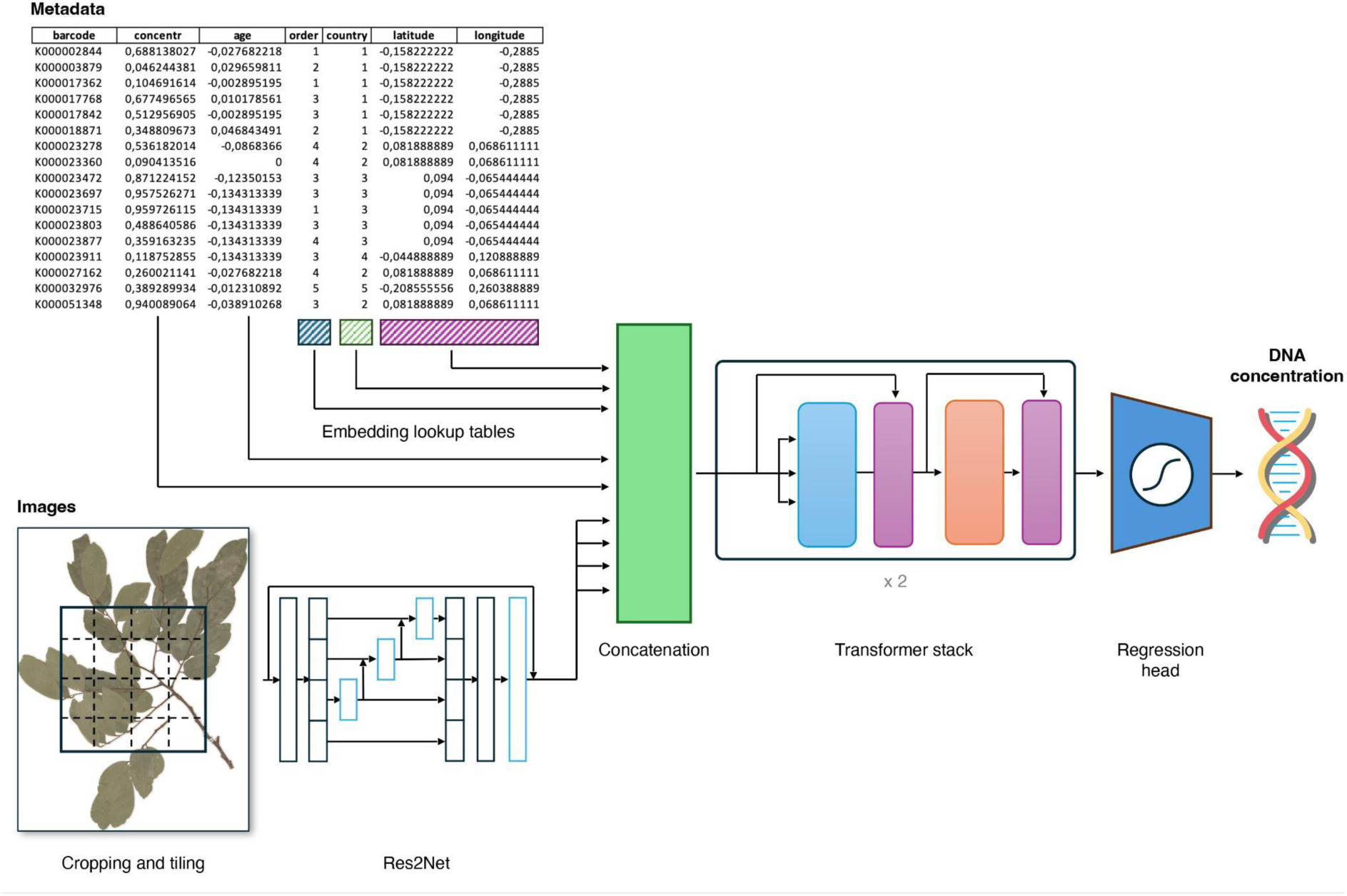
Architecture of the neural network for predicting DNA yield from herbarium samples. The pipeline integrates metadata embedding, image feature extraction via Res2Net, and a transformer stack for processing concatenated features. The final regression head predicts DNA concentration, enabling an accurate assessment of sample quality for genomic studies.

Metadata were processed in parallel. Categorical variables, such as taxonomic order and country of origin, were embedded via learned vectors, with distinct embeddings for missing values. Continuous variables, including the log-transformed sample age and latitude– longitude coordinates, were linearly projected to match the token size used in subsequent network layers.

Image and metadata-derived embeddings are concatenated and augmented with an artificial classification token (CLS). The combined representation is passed through a transformer stack consisting of 2 layers, each containing 4 attention heads and 512-dimensional feedforward layers. Finally, the regression head operates on the CLS embedding after it exits the transformer. That is transformed by two fully connected layers separated by a ReLU activation function, followed by a sigmoid activation mapping predictions to the [0, 1] range. This output corresponds to the quantile-transformed DNA concentration.

### Training Strategy

We initially split the available samples into three distinct sets: training, validation, and testing, with 60%, 20%, and 20% of the samples, respectively.

For hyperparameter tuning, we utilized the Ray Tune library (arXiv:1807.05118), specifically employing the Optuna search algorithm and the Async Successive Halving scheduling strategy (arXiv:1810.05934). The tuning process involved adjusting key parameters, including the number of transformer layers and attention heads, the dimensions of the corresponding feedforward layers, and the dropout rate. Additionally, we optimized the size of the linear layer within the classification head. We set a maximum of 16 epochs for each trial, as we observed that model performance did not improve significantly beyond this point. Trials were terminated early by the search algorithm if no improvement was detected, or they were allowed to run for the full 16 epochs if performance continued to increase.

## Results

### Data Preparation and Cleaning

A deep learning method was developed to predict herbarium-sample DNA concentration using image data and associated metadata. The first stage of the workflow centred on data cleaning, where an OCRNet-based CNN segmented each herbarium sheet to isolate plant material and remove annotations, labels, stamps, and envelopes. After segmentation, a composite dataset was formed by pairing each cleaned image with its pre-processed metadata (taxonomic classification at the order level, latitude–longitude coordinates, and log-transformed sample age), with DNA concentration set as the regression target. Our final dataset comprised 2,029 samples (Table S1).

Mean intersection over union (mIoU) was used to assess segmentation quality on the independent test set of 40 images. After 200 training epochs with data augmentation, the segmentation model achieved an mIoU of 0.9529, indicating a high degree of accuracy in separating plant material from extraneous artifacts.

### Hybrid Neural Network for Regression

Once segmentation was complete and each image–metadata pair was assembled, a hybrid neural network was constructed to integrate both data modalities. Each image was processed by a Res2Net CNN to extract image features. Metadata encoding used learned embeddings for categorical variables and linear projections for continuous variables, creating a unified format for the transformer architecture.

The embeddings were concatenated into token sequences and processed by a transformer stack, where the CLS token captured the final representation. The regression head produced predictions in the [0,1] range, corresponding to quantile-transformed DNA concentrations. Model performance was evaluated on the testing set by comparing predicted DNA concentrations with recorded values. The Pearson correlation coefficient of 0.59 indicates a moderate positive relationship, suggesting that the model captures relevant patterns in the data but also highlighting room for improvement in predictive accuracy.

The results of the training process demonstrated a notable relationship between the quality of color calibration in the input images and the model’s performance. Specifically, we assessed the impact of color jitter augmentation by introducing random variations in hue with a factor of 0.1. This adjustment led to a reduction in the Pearson correlation between the model’s predictions and the ground truth, decreasing from 0.59 to 0.52.

### Sample Classification

Our primary objective was to predict whether a sample has enough DNA for library preparation and sequencing. We selected a minimum of 20 ng/μL as the acceptable DNA concentration, based on the manufacturer’s guidelines (New England Biolabs, n.d.; University of Edinburgh, n.d.; University of California, Davis, n.d).

To evaluate our model’s performance at the classification task, we first used it to predict the DNA concentration of each sample within the testing set. We then applied a threshold to the predictions to obtain a binary labelling. By varying the threshold, we were able to assess the model’s performance across different levels of sensitivity and specificity.

More specifically, for each threshold, we calculated the true positives (samples correctly identified as having sufficient DNA) and false positives (samples incorrectly identified as having sufficient DNA). We then constructed a receiver operating characteristic (ROC) curve (Figure 3) and we obtained a corresponding area under the curve (AUROC) of 0.8, indicating the reliability of our approach in capturing relevant biological patterns. A key advantage of this approach is that it allows the method’s user to tune the model to their desired level of false positive rate (FPR). This flexibility is particularly valuable in applications where sample preservation is crucial, such as historical sample analysis. For example, by aiming at an FPR level of 0.1, our method manages to achieve an accuracy of 0.598, allowing users to prioritize high-confidence sample processing. Ultimately, the user can adjust the trade-off between sensitivity and specificity to suit their specific needs.

**Figure 3.**
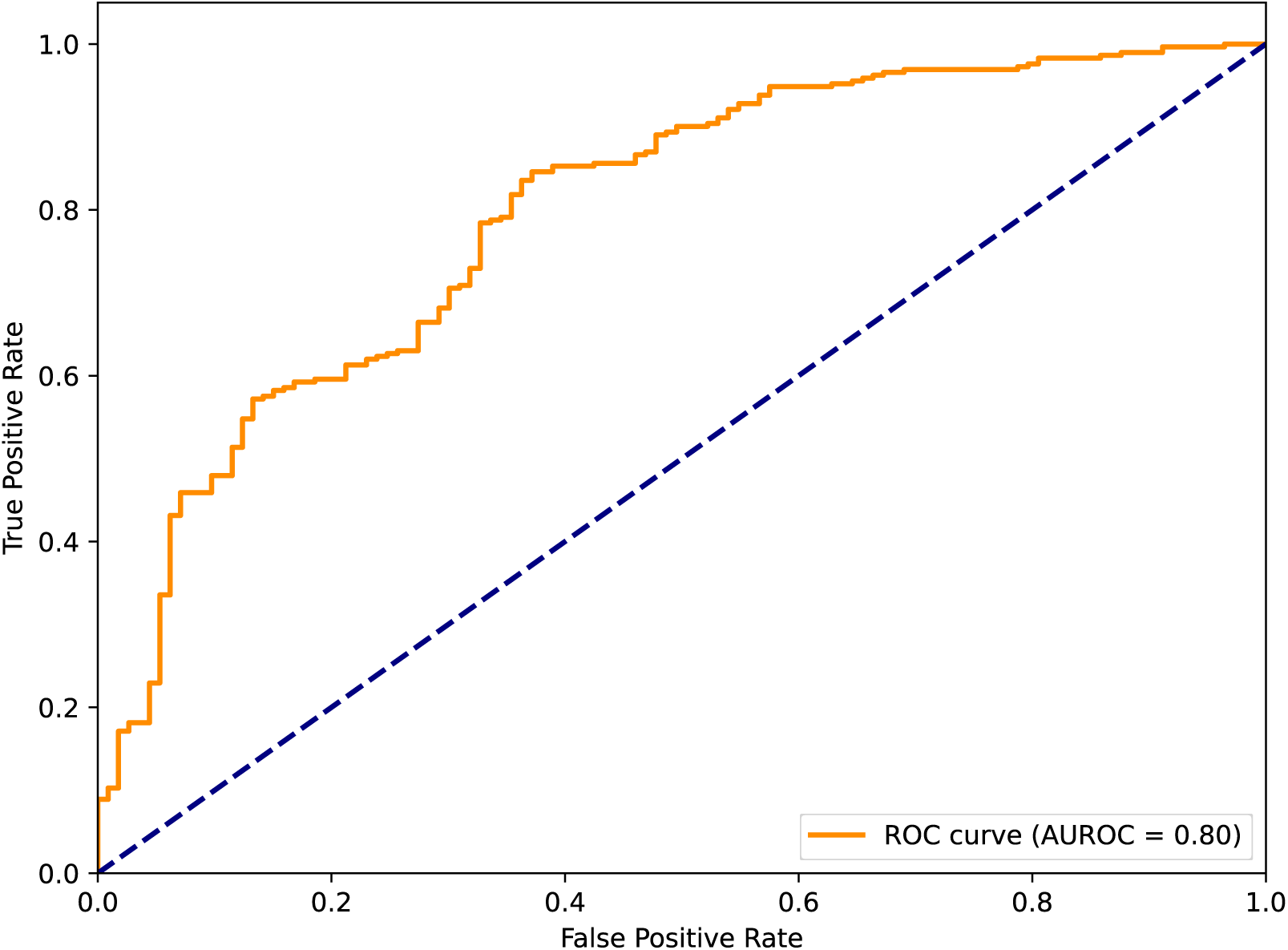
Receiver operating characteristic (ROC) curve illustrating the classification performance, with true positive rate (sensitivity) plotted against false positive rate (1 - specificity) at various classification thresholds.

## Discussion

Herbarium specimens, while valuable for genomic research (Burbano & Gutaker, 2023), present inherent challenges. Many may yield little to no DNA, resulting in the loss of irreplaceable biological material (Marinček et al., 2022). Inefficient or excessive sampling compounds this risk, making informed prioritization crucial for preserving these non-renewable resources. Targeted sequencing approaches, such as Angiosperms353, have demonstrated success in recovering nuclear genes from herbarium specimens, even those over two centuries old, although factors such as climate, collection strategies, and taxon-specific traits significantly influence DNA recovery efficiency (Brewer et al., 2019). Our study addresses these challenges by integrating machine learning with digitized herbarium images and metadata, offering an efficient approach to predicting DNA yield, an important factor that contributes to sequencing success (Vatanparast et al., 2018; Grealy et al., 2019; Marinček et al., 2022). By predicting the likelihood of DNA recovery from herbarium specimens, our approach seeks to optimize sample selection, conserve resources, and minimize damage to collections, in line with recent recommendations (Davis et al, 2024). This strategy aims to improve the usability of herbarium specimens while preserving them for future research.

### DNA yield as a proxy for sequencing success

DNA concentration, though occasionally critiqued as an imperfect proxy for sequencing success (Kates et al., 2021), remains a widely used predictor, particularly in resource-constrained contexts (Vatanparast et al., 2018; Grealy et al., 2019). Low DNA yields restrict genetic analyses to low-depth sequencing or high-copy regions (Ferrari et al., 2023), while relying on total sequencing reads introduces variability due to differences in experimental protocols and re-sequencing efforts (Benjelloun et al., 2020; Illumina, 2014).

Although the number of sequencing reads might appear to be a viable alternative indicator, it is highly dependent on the quality of DNA quantitation (Gua et al., 2013). Furthermore, total read counts often reflect cumulative results from multiple sequencing cycles, potentially misrepresenting the success of the initial sequencing attempt. This can skew predictions and lead to inefficient resource allocation.

We chose to focus on DNA concentration not only for its practicality but also for its technical advantages. Unlike sequencing read yield, which is heavily influenced by external factors such as sequencing depth, budget, and experimental protocols, DNA concentration reflects intrinsic sample quality. Variability in factors like multiplexing, DNA imbalances, and GC content can further complicate the interpretation of sequencing read yield (ThermoFisher Scientific, 2019; Robles et al., 2019; McIntyre et al., 2011). By prioritizing DNA concentration, we isolate the effects of sample quality on sequencing outcomes, offering a more consistent and dependable metric for predicting success.

This focus ensures efficient resource utilization and minimizes the risk of failed sequencing, particularly when resources are limited and the preservation of non-renewable herbarium material is critical (Promega, 2015; Chung et al., 2016; Davis et al., 2024; Illumina, 2025).

### Extending traditional regression-based approaches

Recent advancements in herbarium genomic research, such as the study by Kates et al. (2021), have focused on optimizing specimen selection through regression-based methods. Analysing nearly 8,000 specimens with multivariate linear mixed regression models, Kates et al. identified key predictors of DNA yield and sequencing success, including taxonomic group, source herbarium, and greenness. Greenness emerged as the strongest predictor of DNA yield, while specimen age significantly influenced sequencing outcomes.

Building on these findings, our study employs advanced AI techniques to improve DNA yield prediction. By integrating machine learning with digitized herbarium images and comprehensive metadata, we use convolutional neural networks for effective image segmentation. Our hybrid neural network incorporates a broad range of factors into a unified framework. Unlike regression-based methods (PourMohammad et al., 2020; Mandal & Bhat, 2021), our AI-driven model captures complex, non-linear relationships among predictors, enabling a more holistic and accurate prediction of DNA yield that traditional approaches may overlook.

### Model performance and architectural tradeoffs

Mean intersection over union (mIoU) is a standard evaluation metric for image segmentation tasks. It measures the overlap between the predicted segmentation and the ground truth by computing the ratio of the intersection to the union of these regions, averaged over all classes. Essentially, mIoU quantifies how well the model delineates the areas of interest—in this case, the plant material—from the background and any non-plant elements.

In our study, mIoU was applied to an independent test set of 40 images to evaluate the quality of segmentation. After 200 training epochs with data augmentation, the model achieved an mIoU of 0.9529. This high score indicates that the model has an excellent ability to correctly identify and separate the plant tissue from extraneous artifacts, with very little error. A value of 0.9529, on a scale where 1 represents perfect overlap, confirms that the segmentation performance is outstanding, providing confidence that the extracted plant regions are highly accurate. This represents a significant improvement over traditional image processing techniques, which often struggled to distinguish specimens from labels and other artifacts (Mata-Montero & Carranza-Rojas, 2016; Carranza-Rojas et al., 2017).

The multi-modal architecture we selected for the regression task addressed several important trade-offs. Notably, the transformer architecture can be used to handle missing data, an inherent challenge in herbarium datasets (Heberling, 2021; Eckert et al., 2024; Nicolson et al., 2018). With its built-in capability to handle missing data, this method removes the need for extra preprocessing or imputation steps, which can introduce errors and biases. Furthermore, optimizing the neural network backbone size was crucial for maximizing performance. Increasing the network size did not yield improved results, likely due to the limited size of the dataset. For instance, replacing the Res2Net backbone with 25.7 million parameters (res2net50d.in1k) with a variant containing 45.2 million parameters (res2net101d.in1k) resulted in a decrease in Pearson correlation between predictions and ground truth values, from 0.59 to 0.55.

The relationship between image color calibration and model performance was evident in our results. Applying color jitter, specifically varying the hue by 0.1, caused a decrease in the Pearson correlation from 0.59 to 0.52. This finding suggests that maintaining precise color calibration is crucial for optimizing the model’s predictive accuracy, as even small variations in color accuracy can significantly impact performance. While image capture protocols were beyond the scope of this study, we emphasize the crucial role of maintaining consistent colour calibration across all images. This factor appears to be central to achieving reliable model performance. Future research in this domain should prioritize standardized colour calibration methods during data collection to ensure the model’s generalizability.

As the training dataset grows, the model gains the ability to identify increasingly intricate patterns and relationships within the digitized herbarium images and metadata. This enhanced capacity will eventually allow researchers to input the available specimens for a genomic project and receive a prioritized ranking based on their suitability for DNA sequencing. In the future, this functionality will streamline genomic research by enabling more informed specimen selection, significantly reducing time and resource investment.

### Caveats

While this study represents a significant step forward in optimizing research in herbarium genomics, it is not without limitations. The dataset used to train our machine learning model, while extensive, is inherently constrained by the availability and quality of existing herbarium specimens and their associated metadata. Variability in DNA preservation, collection practices, and storage conditions introduces noise that may affect the potential to generalise our predictions. Additionally, the scope of the study is limited to the specific taxonomic and environmental contexts represented within the dataset, leaving certain plant groups and ecological regions underexplored. Due to the dataset being limited to herbarium samples from the taxonomic groups included in PAFTOL, it inevitably represented only a small subset of the broader scope, as herbarium data comprised just a portion of the overall dataset. With the continued expansion of herbarium genomics research, more data and a broader representation of taxonomic groups will become available, enabling the model to be trained more comprehensively and equitably.

Another limitation of our study is the reliance on a single plant family (ferns) for training the image segmentation model. Our image segmentation model was based on an algorithm that was initially trained using ferns due to their consistent features and availability in curated datasets, providing a solid foundation for developing the algorithm. It was later fine-tuned with angiosperm images, proving highly effective for our dataset and adaptable across diverse plant groups. While this approach yielded high accuracy, it may not fully capture the structural diversity of herbarium specimens from different plant families, which may exhibit varying patterns of DNA degradation and morphological characteristics. Expanding the dataset to include a broader range of species would provide a more generalized model, enhancing its applicability across diverse taxa. Refining the neural network architecture could further improve the model’s ability to handle more complex, multi-dimensional data, thereby enabling the prediction of DNA quality in a wider array of herbarium specimens.

These limitations highlight the need for further data production and collaborative research to enhance the robustness of our approach. Expanding the dataset to include a broader range of species, geographic regions, and storage conditions will improve the model’s predictive power and applicability across diverse research contexts. Moreover, integrating emerging technologies, such as advanced imaging techniques and multi-omics data, could provide deeper insights into the factors influencing sequencing success and further refine specimen selection criteria.

### Future directions for AI in herbarium genomics

The integration of machine learning into herbarium research has unlocked transformative opportunities, enabling large-scale data analysis and expanding the scope of genomic studies. However, the reliability of AI tools remains closely tied to the quality and completeness of image and metadata inputs. Future efforts should focus on incorporating additional variables, such as storage conditions and environmental data, to further enhance predictive power (Hofreiter et al., 2015; Pääbo et al., 2004). This is particularly timely given the proliferation of digitized herbarium collections from major initiatives around the globe.

Several large-scale projects are driving the digitization of herbarium specimens and producing vast datasets suitable for AI applications. For instance, the PAFTOL project at the Royal Botanic Gardens, Kew, aims to sequence the DNA of at least one representative species from all genera of angiosperms, yielding molecular data with an extensive associated database of digitized herbarium images. Similarly, the National Biodiversity Future Center (NBFC) in Italy is generating a significant repository of digitized herbarium specimens as part of its broader biodiversity research framework. Initiatives such as iDigBio (Integrated Digitized Biocollections) in the United States and the Global Biodiversity Information Facility (GBIF) are also providing researchers with unprecedented access to digitized specimen images and metadata from herbaria worldwide.

While resources for digitization and data availability appear less of a limitation than in the past, the irreplaceable and non-renewable nature of herbarium specimens demands careful prioritization to ensure their sustainable use. Advanced AI tools offer a critical opportunity to achieve this by streamlining sample selection, minimizing destructive sampling, and focusing efforts on specimens most likely to yield high-quality genomic data. Learning to fully harness the potential of these irreplaceable resources is essential to avoid<u>ing</u> inefficiencies and wasted opportunities, particularly as the scale of digitized collections continues to grow.

Emerging technologies, such as hyperspectral imaging and representation learning, could further augment these datasets and improve the capabilities of AI-driven tools. For example, Walker et al. (2022) demonstrated how neural networks trained on millions of specimen images yielded transferable insights across diverse research tasks. Applied to herbarium genomics, such technologies will enable researchers to overcome challenges like uneven digitization progress and limited computational resources. Expanding these approaches will facilitate the development of more robust and scalable tools, ensuring herbaria remain invaluable resources for biodiversity research.

As digitization efforts continue to scale, the integration of AI into herbarium genomics is poised to redefine modern biodiversity research. By unlocking new avenues to explore evolutionary patterns, genetic diversity, and species’ adaptive responses to environmental change, AI is poised to support the scientific community in ensuring the optimal use of herbarium specimens while safeguarding them for future generations.

## Data availability

The source code for the model is available on GitHub at the following URL: https://github.com/sales-lab/powerplant

The complete dataset that was meant to be used for model training (before curation) is available upon request.

## Acknowledgements

We sincerely thank the Digital Collections Team at the Royal Botanic Gardens, Kew, for their invaluable support. We are especially grateful to the curators for providing access to the material, the digitizers for imaging, the QA team for uploading the images, Tony Cheuk for facilitating the downloads, and Matthew Clark for assisting with the SFTP access. Their contributions were essential to this work. We would also like to extend our sincere appreciation to Dr. Rhian Smith for her invaluable assistance with the scientific editing of this work. We acknowledge the computing resources utilized to train the model in this study, which were supported by the University of Padova Strategic Research Infrastructure Grant 2017: “CAPRI: Calcolo ad Alte Prestazioni per la Ricerca e l’Innovazione”. The PAFTOL project was funded by grants from the Calleva Project to the Royal Botanic Gardens, Kew. AA acknowledges financial support from the Swedish Research Council (2019-05191; 2024-04303), the Swedish Foundation for Strategic Environmental Research MISTRA (Project BioPath) and RBG Kew Development.

## Author contributions

Yasaman Ranjbaran: design of the research, data collection, writing the manuscript

Olivier Maurin: data collection

Elena Canadelli: data collection

Tomas Morosinotto: data collection

Marie-Helen Weech: data collection

Paul Kersey: data collection

Alexandre Antonelli: design of the research, data collection

William J. Baker: data collection

Gabriele Sales: performance of the research, data analysis, data interpretation, writing the manuscript

Francesco Dal Grande: design of the research, data collection

All authors commented on and edited draft versions of the manuscript.

## Box.1 Methodology of CNN in a nutshell

The methodology used in this research integrates herbarium sheet images and specimen metadata into a unified machine learning framework to predict DNA concentration, addressing a critical need to optimize sequencing efforts for these non-renewable resources. A convolutional neural network (CNN) extracts visual features from the images, identifying key indicators of DNA preservation such as leaf texture, pigmentation, and structural integrity. Metadata, including taxonomic classification (extended to clade and order), geographic origin, and sample age (log-transformed), is transformed into a numerical format for integration.

The extracted visual features and metadata are analyzed together using a transformer neural network, which excels at identifying complex correlations across modalities. By jointly processing these data types, the model captures relationships that would otherwise be missed. For example, leaf texture or pigmentation patterns visible in the images may correlate with environmental factors documented in the metadata, such as collection location or species-specific traits, enabling the discovery of nuanced predictors of DNA concentration.

The hybrid design leverages the synergy between visual and contextual features, capturing correlations that would be lost if processed independently. This integrated approach ensures a more accurate prediction by linking graphical patterns, such as leaf texture or colour variations, to species-specific or environmental characteristics. Such advancements minimize the risks and costs of sequencing, offering a novel, efficient tool for herbarium research. The inclusion of a regression head allows for refined DNA concentration predictions, while optimization techniques enhance model performance, ensuring robustness across diverse taxa and conditions.

